# Hybrid cancer stem cells utilise vascular tracks for collective streaming invasion in a metastasis-on-a-chip device

**DOI:** 10.1101/2024.01.02.573897

**Authors:** Alice Scemama, Sophia Lunetto, Artysha Tailor, Stefania Di Cio, Leah Ambler, Abigail Coetzee, Hannah Cottom, Syed Ali Khurram, Julien Gautrot, Adrian Biddle

**Author notes:** Corresponding author: Adrian Biddle, Centre for Cell Biology and Cutaneous Research, Blizard Institute, 4 Newark Street, London, E1 2AT, UK.

## Abstract

Cancer stem cells (CSCs) drive cancer metastatic dissemination. They do not do so in a vacuum, and the important influence of the tumour microenvironment (TME) on metastatic dissemination is becoming increasingly recognised. Therapeutic targeting of CSC-TME interactions may be a promising route to suppression of tumour metastasis. However, we must first understand how interactions with the TME influence CSC metastatic dissemination. To achieve this understanding, there is a need for experimental models that enable the analysis of dynamic interactions at single cell resolution within a complex environment. To this end, we utilise a metastasis-on-a-chip device to produce a 3D *in vitro* model of CSC interaction with a developing microvasculature, that is amenable to precise imaging and real time studies at single cell resolution. We show that the invasive phenotype of oral squamous cell carcinoma (OSCC) cells is markedly altered when in proximity to a microvasculature, with a switch to a hybrid CSC phenotype that undergoes collective streaming invasion. Mechanistically, ECM compression by the developing vasculature creates an environment that is refractory to cancer invasion, whilst leaving abandoned vascular tracks that can be utilised by hybrid CSCs for collective streaming invasion. Human tissue studies identify streaming invasion in association with vascularised regions in OSCC specimens. These findings elucidate the influence of the vasculature on CSC metastatic dissemination in OSCC, and the role of hybrid CSC invasion plasticity in overcoming this TME constraint.

## Introduction

Tumour metastasis, which seeds secondary tumours, causes the majority of cancer deaths (Mehlen and Puisieux, 2006). Cancer stem cells (CSCs), the sub-population of tumour cells that possess tumour-initiating potential (Clarke et al., 2006, Driessens et al., 2012), can adopt phenotypes that drive tumour invasion and metastasis (Biddle et al., 2011, Lawson et al., 2015, Pastushenko et al., 2021, Youssef et al., 2023) and express heightened resistance to therapy (Biddle et al., 2016, Kreso et al., 2013). Epithelial-to-mesenchymal transition (EMT), a developmental process in which epithelial cells acquire a migratory mesenchymal phenotype, is re-activated by CSCs to drive tumour invasion and migration to secondary sites (Yang et al., 2008). Mesenchymal-to-epithelial transition (MET), where migratory CSCs revert back to an epithelial phenotype, enables new tumour growth at metastatic sites (Tsai et al., 2012). The ability to undergo these phenotypic changes is referred to as plasticity, and is a defining feature of CSCs (Chaffer et al., 2016). However, the role of interactions with the tumour microenvironment (TME) in shaping CSC plasticity is not well understood. Such interactions may act to either enhance or repress tumour metastasis, and may present promising therapeutic targets (Ferguson et al., 2021).

The TME consists of the cellular and extracellular matrix (ECM) components that surround the tumour and make up the peri-tumoural stroma. These include immune cells, fibroblasts and a vasculature, all contained within an ECM hydrogel (Anderson and Simon, 2020). The angiogenic tumour vasculature is a key component of the TME, providing oxygen to the tumour and removing waste products. It also serves as the route for circulating immune cells to access the tumour, and as a major route for tumour metastatic dissemination. However, it is not known whether interaction with the developing vasculature can actually play a role in shaping CSC phenotype. CSCs have been observed to concentrate within a vascular niche in OSCC xenografts (Krishnamurthy et al., 2010). A TGFβ-driven vascular niche signalling network has been shown to maintain CSCs in a genetically engineered mouse model of cutaneous SCC (Oshimori et al., 2015). However, the latter was due to signalling between macrophages and cancer cells, rather than a direct role of the vasculature (Taniguchi et al., 2020). Thus, it is still not known whether the vasculature plays a direct role in shaping CSC phenotype. To investigate this, an experimental model where this relationship can be tested in isolation is required; the complex TME of a mouse model is not a suitable experimental system.

Advances in human tissue analysis has enabled characterisation of metastatic CSCs within the complex human TME (Jensen et al., 2015, Youssef et al., 2023). However, these can only present a static picture and are unable to model dynamic interactions. They also preclude mechanistic studies. CSC research therefore relies primarily upon mouse models and simple cell culture models, but these are also not well suited to the analysis of dynamic interactions with a complex TME. Simple cell culture models lack cellular complexity and a 3D environment. Mouse models possess a complex environment, but this is not a human tumour environment and working with live animals presents formidable challenges for detailed analysis of cellular interactions. The incorporation of cellular complexity within macro-scale 3D *in vitro* models also poses challenges for detailed analysis, with sub-optimal imaging depth and inability to compartmentalise populations. To overcome these challenges, and enable analysis of CSC interactions with a complex TME within a 3D *in vitro* model, we have utlilised a microfluidic experimental platform.

Microfluidic metastasis-on-a-chip models make use of microfluidic designs to generate 3D culture systems. They are typically comprised of a horizontal compartmentalised structure with multiple adjacent microchannels, and can recapitulate the barrier function of the microvasculature and its ability to perfuse molecules and cells (Kim et al., 2013, Lee et al., 2014). Fibrin is commonly used as the ECM component within these devices due to its ability to support a developing vasculature. Microfluidic devices are attractive platforms for the study of tumour metastasis, as the channels enable compartmentalisation of different cell types and ECM components. Composed of a clear polymer attached to a transparent cover slip, microfluidic devices are optimal for precise imaging and real-time studies (Scemama et al., 2022). This enables superior analysis of dynamic interactions at single cell resolution. For example, it has been shown that secretion of TNFα by macrophages causes endothelial layer impairment and consequent promotion of both intravasation and extravasation events in a metastasis-on-a-chip model (Chen et al., 2013, Zervantonakis et al., 2012). However, these models have not yet been used to study the effect of a vasculature on tumour heterogeneity at single cell resolution, including that relating to CSC plasticity. Modulation of EMT markers has been observed within microfluidic systems, but tumour cell populations were treated as homogeneous entities and spatial interactions were not investigated (Lee et al., 2018, Rizvi et al., 2013).

Oral squamous cell carcinoma (OSCC) is one of the top ten cancers worldwide, with over 300,000 cases annually, and an increasing incidence (in the UK, incidence has increased by 23% over the past decade). The majority of cases are human papilloma virus (HPV)-negative. HPV-negative OSCC is a deadly disease with frequent metastatic spread, which is the single most important predictor of outcome (Sano and Myers, 2007). It is also exceptionally amenable to *in vitro* experimental modelling of *in vivo* tumour behaviour (Biddle et al., 2016, Youssef et al., 2023). Here, we investigate the impact of a microvasculature on OSCC CSC plasticity within a microfluidic chip model. We find that proximity to a microvasculature alters the phenotype of invading CSCs and their mode of invasion. Mechanistically, this is due to ECM remodelling by the developing vasculature and co-option of vascular channels by invading CSCs. Analogous changes in tumour phenotype are seen in vascularised regions of human OSCC specimens.

## Results

### Proximity to a microvasculature alters the mode of invasion of OSCC

Having first optimised co-culture in the microfluidic chip and analysis of invasion **(Figure S1)** (Scemama et al., 2023), we proceeded to assess the effect of a microvasculature on the mode of invasion of OSCC cells. We cultured the CA1 and LUC4 OSCC cell lines in the presence and absence of HUVECs in the microfluidic chip. HUVECs were first added, and allowed to grow into the central fibrin gel and form sprouting structures until their growth had reached approximately half way across the 700 µm wide middle channel. OSCC cells were then added in the opposite side channel, in both chips containing a microvasculature and control chips without a microvasculature **(Figure 1A)**. In the presence of a microvasculature, OSCC invasive area within the central channel was reduced **(Figure 1B, C)**. This was accompanied by an increased length of the invasive front perimeter in both OSCC cell lines, but not in the NTERT non-transformed keratinocyte cell line **(Figure 1D)**. In control chips, the OSCC cells invaded as large clusters with a broad front, whereas in vascularised chips there were smaller and thinner invasive clusters **(Figure 1B, white arrows)**. These had the appearance of streams of invasive cells, which create a longer invasive front perimeter. This stream-like phenotype was observed with both the CA1 and LUC4 OSCC cell lines, but not with NTERTs (**Figure 1E**). Altogether, this data suggests that vascular endothelial cells create an environment refractory to OSCC invasion on a broad front, which instead invades as thin streams of cells, a phenotype not observed in a non-transformed cell line.

**Figure 1.**
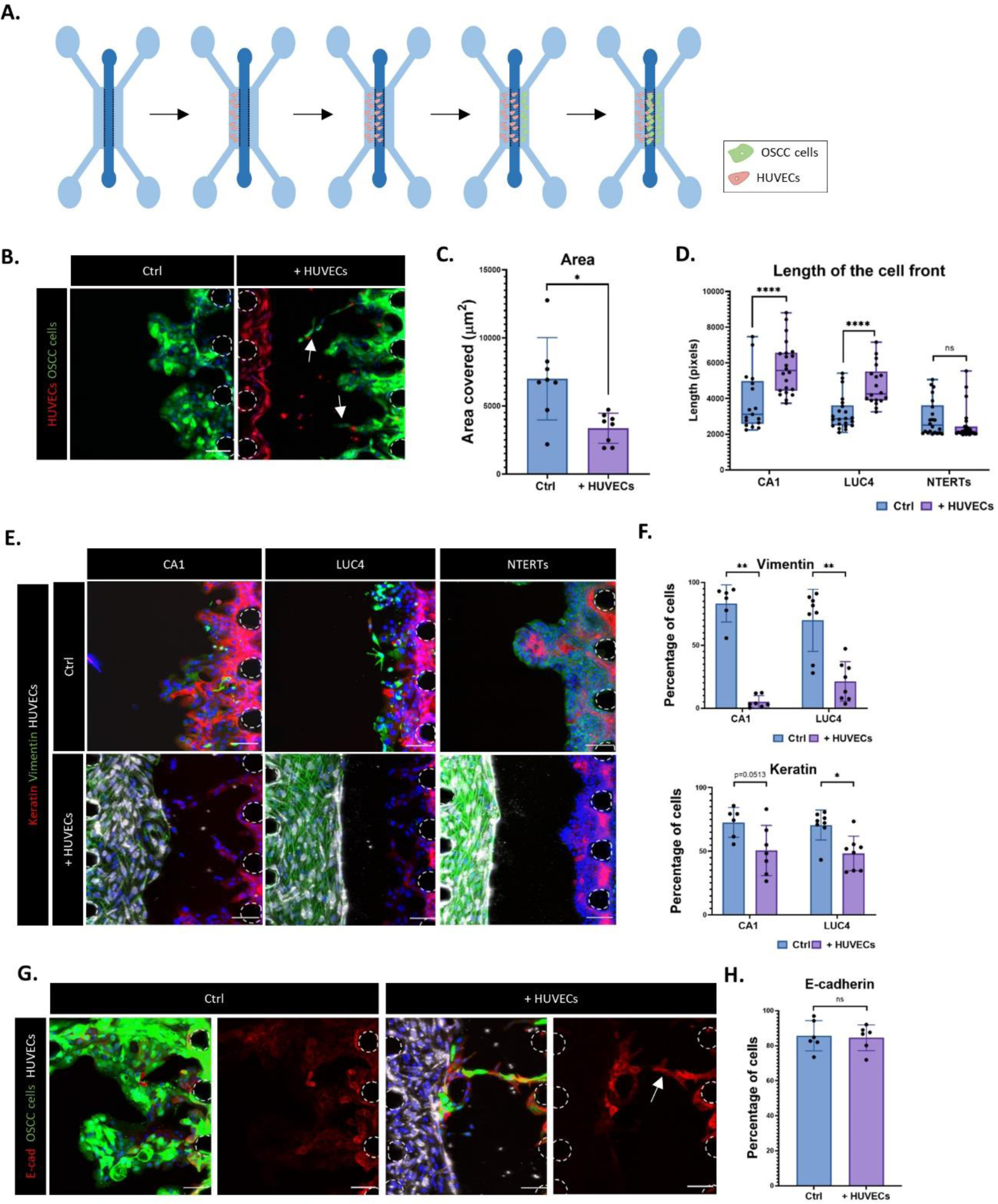
Proximity to a microvasculature alters the mode of invasion of OSCC. **A.** Experimental set-up for assessing the invasion of OSCC cells into a vascularised fibrin gel in the microfluidic chip. HUVECs are added to the left channel on day 1, they vascularise the central gel channel, then OSCC cells are added to the right channel on day 3-4. Cells are fixed and analysed at day 10-11 **B.** HUVECs (Red, RFP) and CA1 OSCC cells (Green, GFP) in the central fibrin gel channel. White arrows indicate streams of invading OSCC cells. **C.** Area covered by the CA1 cells within the central channel. **D.** Total perimeter length of the invasive front for the CA1 and LUC4 OSCC cell lines and NTERT non-transformed keratinocyte line. Mann-Whitney test. Data from 4 independent experiments, with each dot representing a single field of view of a microfluidic chip. **E.** Immunofluorescent staining of cells invading into the central channel, with and without HUVECs. Red, Keratin; Green, Vimentin; Grey, RFP-HUVECs; Blue, DAPI. **F.** Percentage of OSCC cells expressing Vimentin (top) and Keratin (bottom). Mann-Whitney test. **G.** Immunofluorescent staining for E-cadherin in invading CA1 cells. For each condition (+/− HUVECs), the merge is on the left and single channel for E-cadherin on the right. Red, E-cadherin; Green, CA1-GFP; Grey, RFP-HUVECs; Blue, DAPI. The white arrow indicates a stream of invading OSCC cells. **H.** Percentage of CA1 cells expressing E-cadherin. For all microscopy images, posts are highlighted with white dashed lines and scale bars: 100 µm.

Cancer cells can employ distinct modes of invasion during metastasis, including single cell invasion, multicellular streaming invasion and collective invasion (Friedl and Alexander, 2011, Friedl et al., 2012, Pandya et al., 2017). Multicellular streaming has been described as a migratory phenotype where cancer cells migrate one after the other as chains of cells with weak cell-cell adhesion, with the quasi-collective shape driven largely by constraints imposed by the environment (Pandya et al., 2017). On the other hand, collective invasion encompasses different forms, ranging from thin stream-shaped masses to large and wide ones. They have the consistent feature of maintaining cell-cell junctions including adherens junctions (Friedl et al., 2012). We found that E-cadherin, a transmembrane glycoprotein and key component of adherens junctions (van Roy and Berx, 2008), was strongly expressed at the cell surface in the streams of cells **(Figure 1G, H)**. This indicates that the streams of invading OSCC cells maintain cell-cell junctions and therefore appear to be undergoing collective invasion. Epithelial-mesenchymal transition (EMT), marked by the intermediate filament Vimentin (Thiery, 2002), is associated with the non-collective forms of invasion (single cell and multicellular streaming). Vimentin was markedly reduced in the OSCC cells in the vascularised chips, alongside a smaller but significant reduction in Keratins (pan-keratin antibody) **(Figure 1E, F)**. Whilst Vimentin can be seen in individual invading cells at the invasive front in control conditions (particularly in LUC4), it is absent from the invasive streams in the presence of a microvasculature. This indicates an absence of EMT in the invading streams, further supporting the conclusion that they are undergoing collective epithelial invasion.

### Invasive OSCC streams express a hybrid CSC phenotype in vascularised chips

OSCC invasion has been previously associated with a CSC phenotype with CD44^high^EpCAM^low^ marker expression (Biddle et al., 2011), and CD44 is an established marker for OSCC CSCs (Prince et al., 2007). We therefore characterised the expression of CD44 and EpCAM in the invading streams of cells **(Figure 2A-C)**. CD44 expression was restricted to the invasive front in both control and vascularised conditions, but the CD44+ invasive front demonstrated marked changes between conditions. In control chips, the CD44+ invasive front was broad, covering the whole front of the invasive tumour mass. In vascularised chips, the CD44+ invasive front was restricted to the invasive streams **(Figure 2A)**. Quantification of mean invasive distance for cells expressing CD44 and EpCAM demonstrated that CD44+ cells were at the invasive front whereas EpCAM+ cells were towards the rear of the invasive tumour mass, in both control and vascularised chips **(Figure 2B)**. Consistent with this, the cells at the invasive front were CD44^high^EpCAM^low^, and these were restricted to the invasive streams in the vascularised chips **(Figure 2A, white arrows)**.

**Figure 2.**
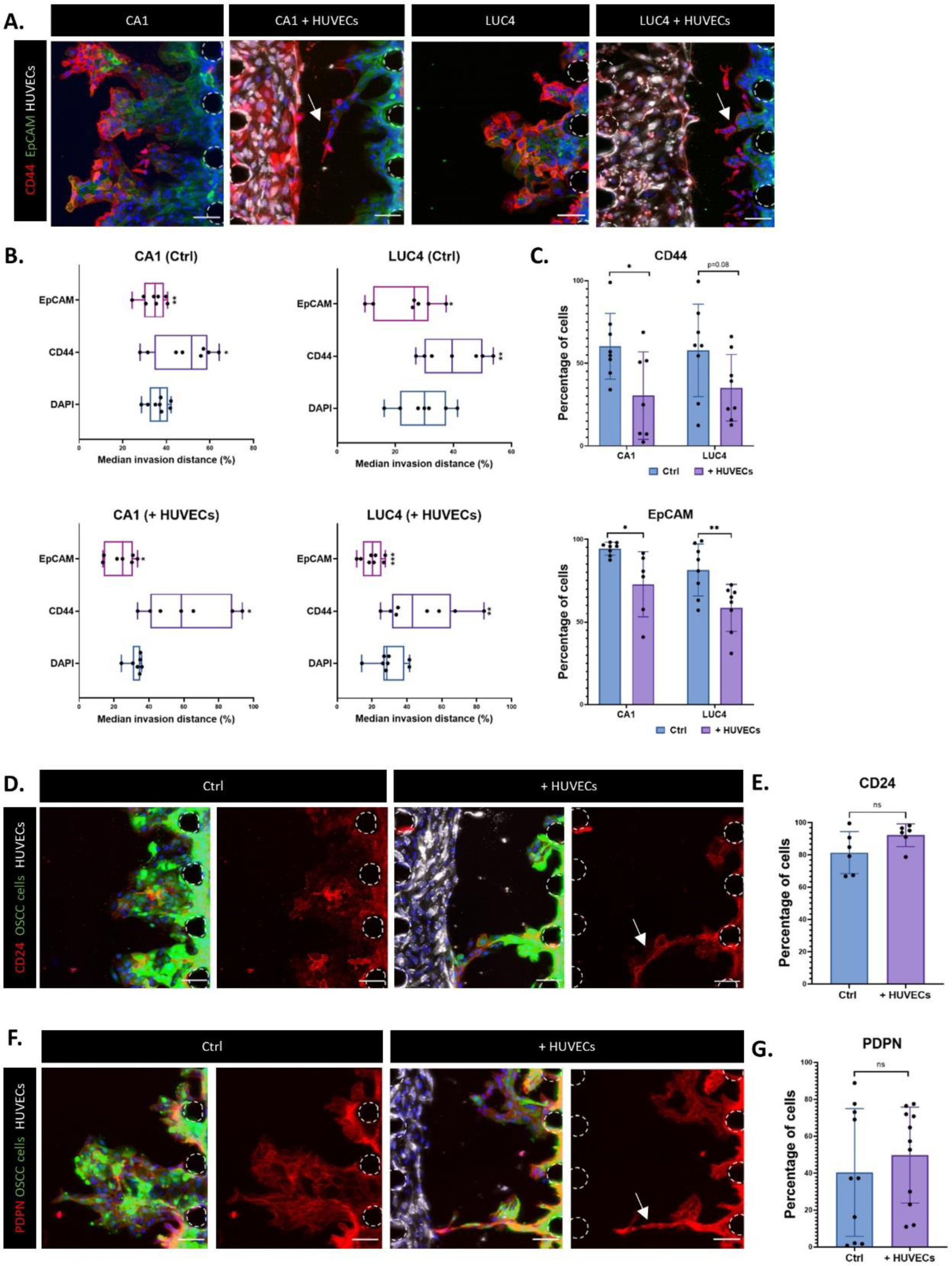
Invasive OSCC streams express a hybrid CSC phenotype in vascularised chips. **A.** Immunofluorescent staining for CD44 and EpCAM in invading CA1 and LUC4 OSCC cell lines. Red, CD44; Green, EpCAM; Grey, RFP-HUVECs; Blue, DAPI. The white arrows indicate streams of invading OSCC cells. **B.** Median invasion distance of OSCC cells positive for CD44, EpCAM, or DAPI (whole population), when cultured alone (top) and with a vasculature (bottom). Invasive distance into the central channel was normalised to the maximum invasion distance for the whole OSCC cell population. Paired t-test. **C.** Percentage of CA1 and LUC4 cells expressing CD44 (top) and EpCAM (bottom). Mann-Whitney test. **D, F.** Immunofluorescent staining for CD24 **(D)** and PDPN **(F)** in invading CA1 cells. For each condition (+/− HUVECs), the merge is on the left and single channel for CD24/PDPN on the right. Red, CD24/PDPN; Green, CA1-GFP; Grey, RFP-HUVECs; Blue, DAPI. The white arrows indicate streams of invading OSCC cells. **E, G.** Percentage of CA1 cells expressing CD24 **(E)** and PDPN **(F)**. For all microscopy images, posts are highlighted with white dashed lines and scale bars: 100 µm.

Invasive CD44^high^EpCAM^low^ CSCs in OSCC often undergo EMT (Biddle et al., 2011, Youssef et al., 2023), but the lack of Vimentin and retention of E-cadherin demonstrates an absence of EMT in the invasive streams in the vascularised chips. The observed concurrent downregulation of the epithelial marker EpCAM **(Figure 2A-C)** and epithelial keratins **(Figure 1E, F)** suggests that the OSCC cells invading as streams of cells appear to lose some epithelial traits while not undergoing a full EMT, indicating potential adoption of a hybrid CSC state. Such a state has been associated with metastasis in animal models of squamous cell carcinoma (Pastushenko et al., 2021) and human OSCC specimens (Puram et al., 2017).

CD24 (Biddle et al., 2016) and PDPN (Puram et al., 2017) (A. Biddle, unpublished) have been identified as markers for hybrid CSCs with increased plasticity in OSCC. To further investigate whether the invasive streams are associated with a hybrid CSC phenotype, we assessed the expression of CD24 and PDPN in the chips **(Figure 2D-G)**. Both of these markers were strongly expressed in the invasive streams **(Figure 2D, F, white arrows)**, supporting the conclusion that OSCC cells invading towards a microvasculature form thin collective streams with a hybrid CSC phenotype **(Figure 3A)**.

**Figure 3.**
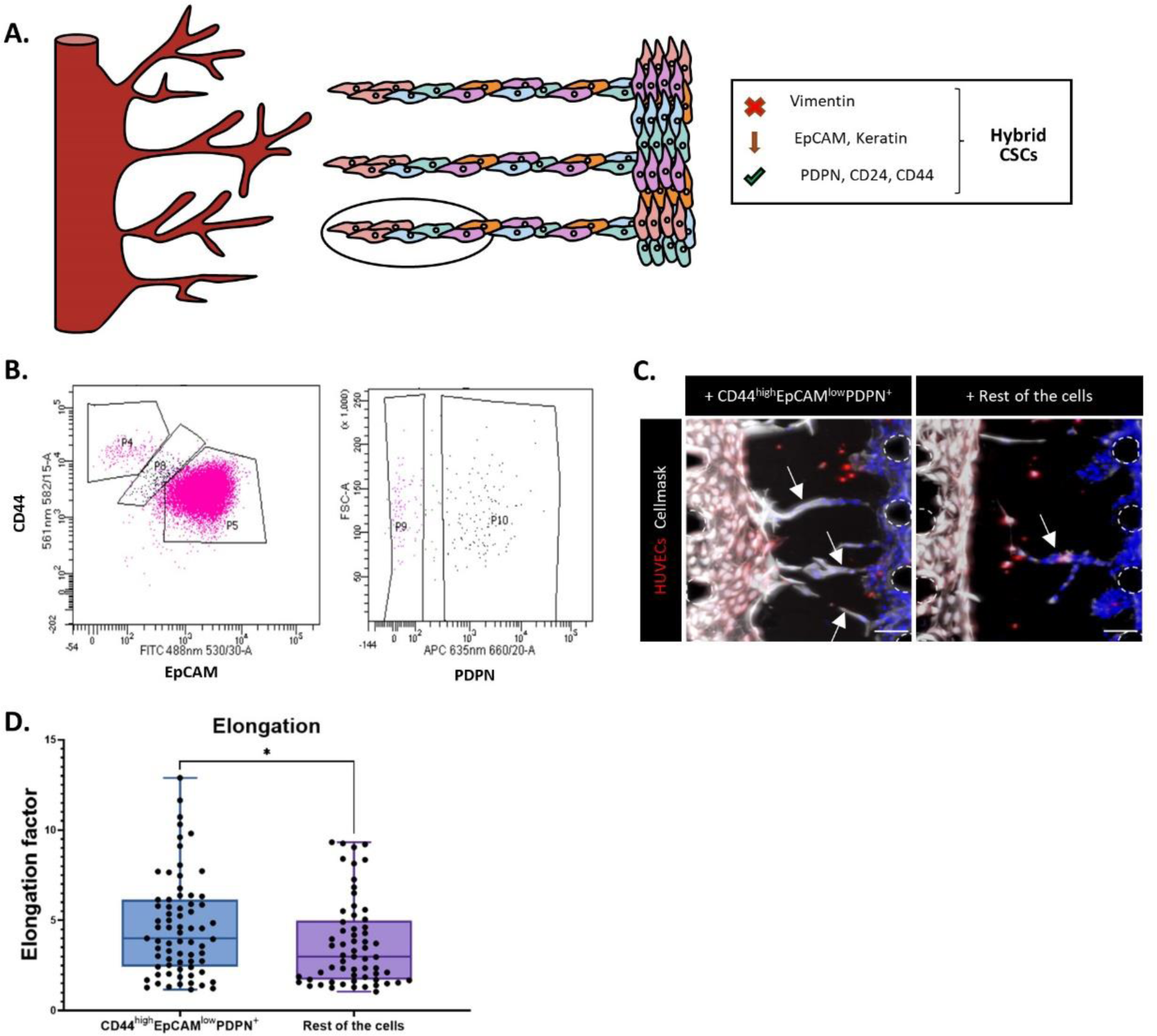
The hybrid CSCs that form the invasive streams are a pre-existing population in OSCC. **A.** Hybrid CSC markers in OSCC cells undergoing streaming invasion towards a vasculature. There is downregulation of epithelial keratins and the EMT marker Vimentin, alongside expression of the hybrid CSC markers CD44, CD24 and PDPN. This informs the gating strategy for enriching these cells through cell sorting. **B.** The gating strategy for enrichment of CD44^high^EpCAM^low^PDPN^+^ cells from the CA1 OSCC cell line by flow cytometric cell sorting. The EMT population (P4) and majority epithelial population (P5) are excluded. P10 is the PDPN+ derivative of P8. P10 cells were sorted and compared to a sorted population comprising all other live cells. Left plot: CD44 (y-axis) and EpCAM (x-axis). Right plot: Forward scatter (FSC) (y-axis) and PDPN (x-axis). **C.** Sorted CD44^high^EpCAM^low^PDPN^+^ cells (left), and all other live cells (right), invading towards HUVECs in the central channel. Red, RFP-HUVECs; Grey, Cell mask; Blue, DAPI. The white arrows indicate streams of invading OSCC cells. Post are highlighted in white (dashed line). Scale bars: 100 μm. **D**. Elongation factor of the invasive cell masses for the two sorted populations. Mann-Whitney test. Data from 3 independent experiments, with each dot representing a cell mass in a microfluidic chip.

### The hybrid CSCs that form the invasive streams are a pre-existing population in OSCC

To understand whether the cells invading as streams of cells are pre-existing in the parental cell line or emerge *de novo* in the presence of vascular endothelial cells, we isolated cells with the hybrid CSC marker profile from the CA1 OSCC cell line in 2D culture. We isolated the CD44^high^EpCAM^low^PDPN^+^ population and all other live cells from the CA1 cell line by fluorescence-activated cell sorting (FACS) **(Figure 3B),** and compared the behaviour of these two populations in vascularised microfluidic chips **(Figure 3C, D)**. The CD44^high^EpCAM^low^PDPN^+^ population formed an enhanced number of streams, observed both visually **(Figure 3C, white arrows)** and through quantification of the elongation factor of invasive masses **(Figure 3D)**. Therefore, the stream-forming population is pre-existing and has a hybrid CSC marker profile.

### Invasive OSCC streams form within matrix that is remodelled by the developing vasculature

We next investigated the mechanism through which proximity to a microvasculature induces this change in the OSCC invasive phenotype. It has previously been shown that, during angiogenesis, remodelling of the matrix is observed in close proximity to the sprouting vasculature (Juliar et al., 2018, Kniazeva et al., 2012, Rüdiger et al., 2020). Remodelling-induced increases in matrix density can hinder cell invasion (Ehrbar et al., 2011, Zaman et al., 2006), and we therefore assessed whether the developing vasculature drives matrix remodelling in the microfluidic chip. Reflectance imaging to visualise the matrix fibres showed that, in vascularised microfluidic chips, there is remodelling to produce thicker fibrin fibres. This effect is then partially reversed after subsequent addition of the CA1 OSCC cell line **(Figure 4A-C)**. An accumulation of thicker fibrin fibres was observed at the growing front of the sprouting vasculature, suggesting active remodelling of the matrix by the developing vasculature **(Figure 4D)**. When fluorescently-tagged microparticles were incorporated into the fibrin matrix, these were displaced by the developing vasculature, resulting in an accumulation of these near the growing front **(Figure 4E-G)**. Hence, as the vascular endothelial cells grow into the fibrin matrix and form a sprouting vasculature, these cells remodel and compress the matrix leading to the aggregation of fibrin fibres at the front of the developing vasculature. This creates a denser environment, which may be refractory to OSCC cell invasion. In support of this conclusion, artificially increasing matrix density by increasing the fibrinogen concentration also resulted in reduced OSCC invasion and increased invasive front elongation in the presence of a microvasculature **(Figure S2)**.

**Figure 4.**
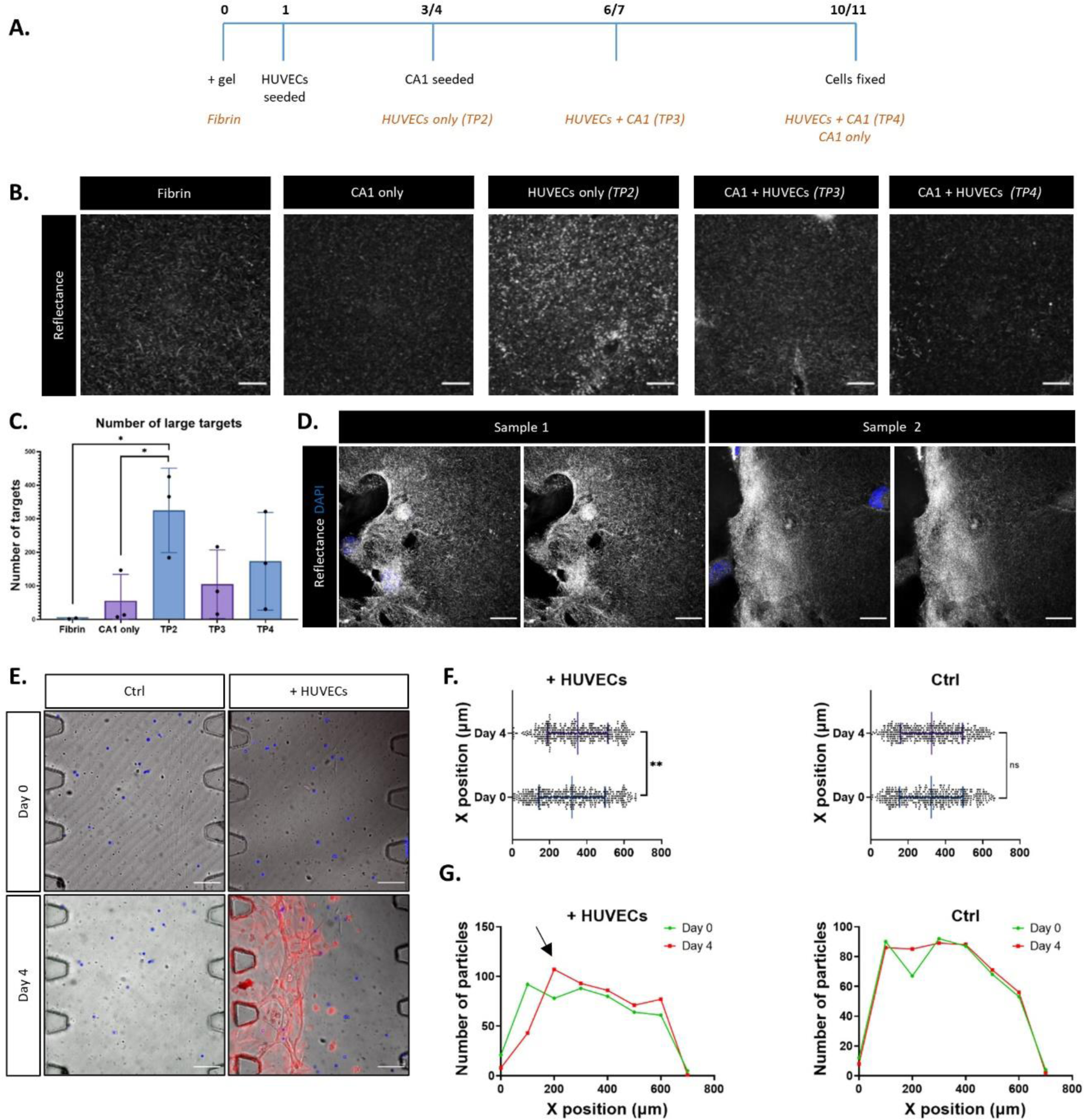
Invasive OSCC streams form within matrix that is remodelled by the developing vasculature. **A.** Experimental timeline. Black text indicates experimental stages, orange text indicates the sampling timepoints for different experimental conditions. **B.** Reflectance imaging of fibrin fibres at the sampling timepoints for different experimental conditions indicated in **(A)**. Grey, Reflectance signal. Scale bars: 20 μm. **C.** Number of large fibrin fibres (large targets) in the reflectance imaging. **D.** Fibrin fibres at the HUVEC cell front at day 3-4. HUVECs are progressing from the left. Grey, Reflectance; Blue, DAPI. For each sample, the composite is on the left and the single reflectance channel is on the right. Scale bars: 20 μm. **E.** Fluorescent particles (blue) and RFP-HUVECs (red) in the central channel at day 0 and day 4. Scale bars: 100 μm. **F.** X position (distance into the central channel measured along a line starting at, and perpendicular to, the left side of the channel) of the fluorescent particles in the control (ctrl) samples and samples with HUVECs. Mann-Whitney test. Data from 3 independent experiments, with each dot representing a fluorescent particle in a single microfluidic chip. **G.** Histograms of the particle number for each X position (20 μm ranges) in the control (ctrl) samples and samples with HUVECs, showing a compression of the particle population (black arrow) near X position zero, which is the point of HUVEC entry.

To investigate whether the effect on OSCC invasion is spatially located to the areas of increased matrix density, we employed new microfluidic chip designs with a greater separation between the HUVEC and OSCC cell seeding points. These included a 4-channel device where two fibrin gels were added in two separate 700 µm wide middle channels, and a microfluidic chip with a 2000 µm wide middle channel (**Figure 5A**). In these settings, no increase in invasive OSCC streams was observed in the vascularised chips **(Figure 5B, C)** and the overall invasive area of the OSCC cells was not reduced compared to the control chips **(Figure 5D)**. Furthermore, accumulation of thicker fibrin fibres was restricted to the vascularised region, being absent from areas closer to the OSCC cell seeding point **(Figure 5E-H)**. Therefore, when OSCC invasion is outside the area of matrix remodelling by the developing vasculature, there is no effect on the mode of invasion. Only OSCC invasion proximal to the developing vasculature, within the region of matrix remodelling, is prevented from invading across a broad front and instead adopts a streaming invasive phenotype.

**Figure 5.**
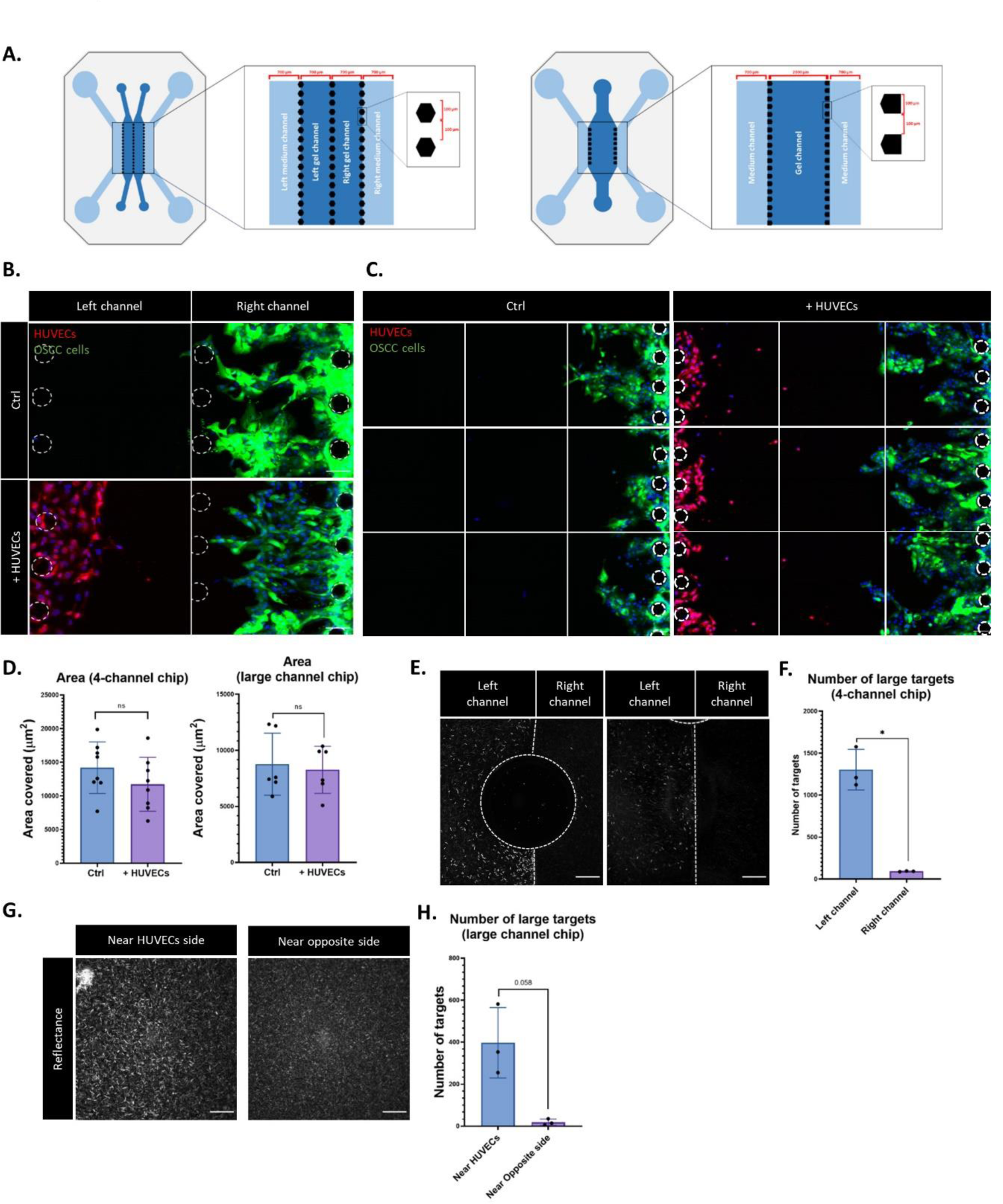
The effect on OSCC invasion is spatially located to the areas of increased matrix density. **A.** Modified chip designs. Left: A 4-channel microfluidic chip, with each channel width measuring 700 μm and separated with a line of posts. Fibrin matrix is added to both central channels. Right: A wider 3-channel microfluidic chip, with the side channels being 700 μm and central channel being 2000 μm wide. **B.** CA1 cells only (ctrl) or with HUVECs in the 4-channel chip, showing the left and right central channels. **C.** CA1 cells only (ctrl) or with HUVECs in the wider 3-channel chip, with stitched fields of view covering the entire width of the 2000 μm central channel. For **(B)** and **(C)**, Green, GFP-CA1; Red, RFP-HUVECs; Blue, DAPI. Posts are highlighted in white (dashed line). Scale bars: 100 μm. **D.** Area covered by the CA1 cells within the central channel. **E.** Reflectance imaging at day 3-4 of fibrin fibres in the left and right central channels of the 4-channel chip with HUVECs only. Two examples are shown, with HUVECs seeded at the left-hand side. White dotted lines mark out the boundary between channels and posts. Grey, Reflectance. Scale bars: 20 μm. **F.** Number of large fibrin fibres (large targets) in the reflectance imaging of the 4-channel chip. **G.** Reflectance imaging at day 3-4 of fibrin fibres at two different locations in the central channel of the wider 3-channel chip with HUVECs only. Grey, Reflectance. Scale bars: 20 μm. **H.** Number of large fibrin fibres (large targets) in the reflectance imaging of the wider 3-channel chip.

### Co-option of angiogenic tracks for OSCC streaming invasion

The creation of a denser matrix environment may explain how the developing vasculature is refractory to OSCC invasion, but it remained unclear why the OSCC cells invade as streams in this environment.

Upon addition of OSCC cells to the chips, we consistently observed regression of the microvasculature **(Figure S3A)**. This was accompanied by reduced proliferation **(Figure S3B)** and reduced dextran permeability **(Figure S3C)**. This effect was not specific to OSCC cells, as we also observed microvasculature regression in the presence of NTERT non-transformed keratinocytes **(Figure S3D)**. Therefore, it may represent a process of vascular maturation in response to crosstalk with epithelial tissues. Another recurring observation from our experiments is that, during vascular development, individual endothelial cells often progressed all the way across the middle channel before retreating and appeared to leave disturbances in the matrix. We hypothesised that these disturbances (possibly a physiological mechanism to guide vascular sprouting) may form tracks through which OSCC streaming invasion can progress in the context of an overall matrix environment that is compressed and less receptive to invasive infiltration.

We therefore investigated whether the regressing microvasculature leaves tracks in the matrix that are utilised for OSCC streaming invasion. Images of the middle channel were taken at multiple experimental timepoints, both before and after the addition of OSCC cells. These showed single endothelial cells producing tracks within the matrix during vascular development, followed by utilisation of these tracks for OSCC streaming invasion **(Figure 6 and Figure S4)**. This provides an explanation for the observed formation of invasive OSCC streams within vascularised chips.

**Figure 6.**
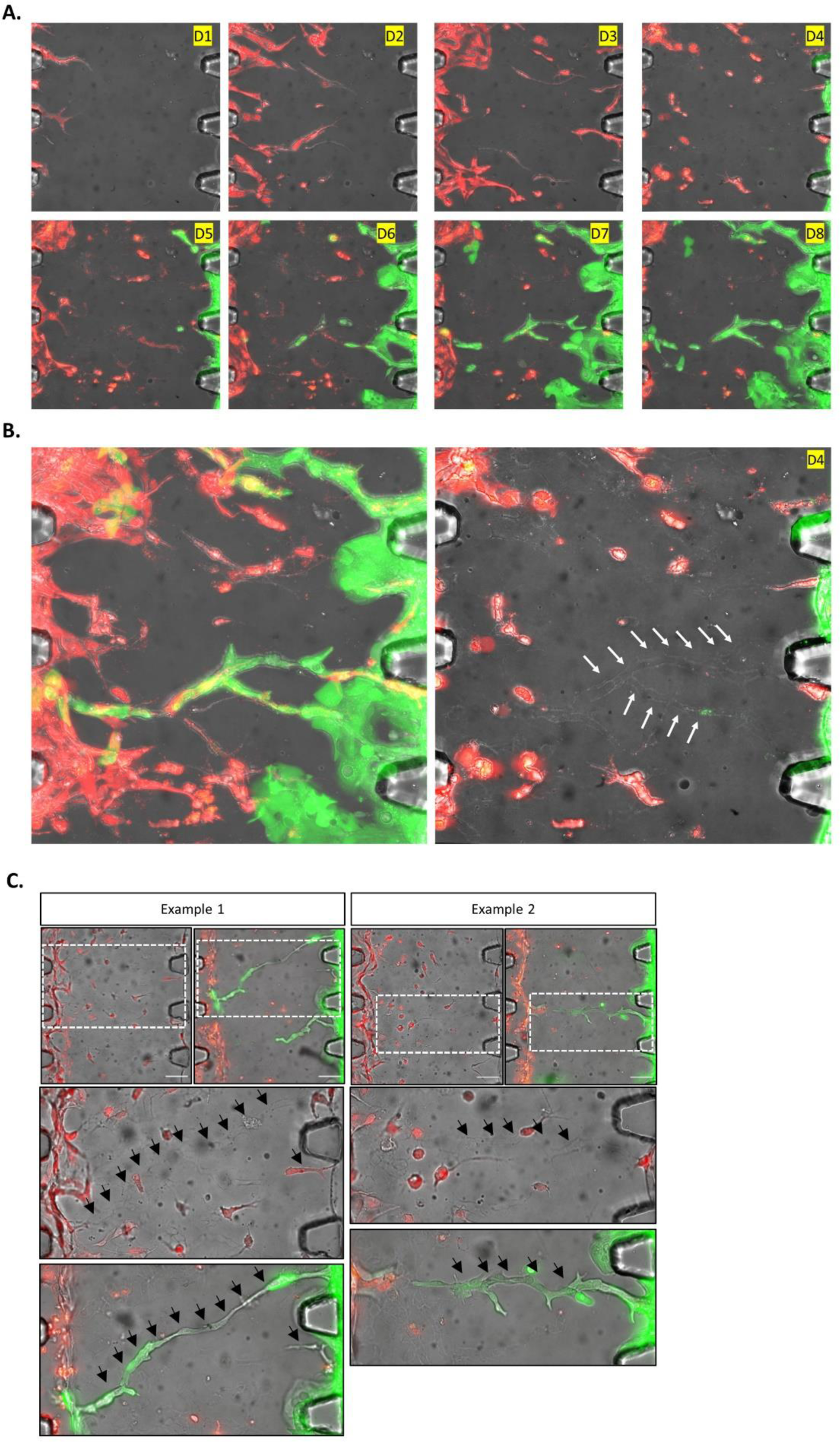
Co-option of angiogenic tracks for OSCC streaming invasion. **A.** Daily timepoint analysis at days 1 to 8 in a vascularised chip, with HUVECs (Red; RFP) added at day 1 (D1) and CA1 OSCC cells (Green; GFP) added at day 4 (D4). **B.** Left: Overlay projection of all timepoints from A, showing the CA1 OSCC cells (Green; GFP) taking the same route through the matrix as the HUVECs (Red; RFP). Right: The day 4 timepoint image with the HUVEC-produced tracks denoted with white arrows. A single RFP+ HUVEC can be seen at the head of a track, having progressed the whole way across the central channel. **C.** Examples of the tracks generated by the HUVECs, showing the same field of view before addition of CA1 OSCC cells (left image, and top inset) and after CA1 addition (right image, and bottom inset). Black arrows denote the tracks produced by the HUVECs (Red; RFP), which are then utilised by the OSCC cells (Green; GFP) for invasion. Single RFP+ HUVECs can be seen at the head of each track before CA1 addition, having progressed the whole way across the central channel. Scale bars: 100 μm.

### Association of streaming invasion with vascularised regions in human OSCC specimens

Invading stream-like projections have previously been associated with poorer prognosis in analysis of H&E-stained sections from human tongue squamous cell carcinoma (Wu et al., 2019), but proximity to vasculature has not been investigated. Therefore, to determine whether the behaviour manifested in our *in vitro* model can be observed *in vivo*, we analysed H&E sections from human OSCC tumours. Given the CSC phenotype and downregulation of epithelial differentiation markers in the invasive streams in our microfluidic chips, we devised a H&E scoring system to account for both differentiation status and streaming phenotype. We assessed the phenotype of cancer cell clusters by classifying them as either 1) well differentiated (containing keratin pearls), 2) less differentiated, or 3) stream-like. A scoring system was used with each keratin pearl scoring −2, less differentiated clusters scoring 1, and stream-like clusters scoring 2. A region scoring low would thus have higher levels of differentiation with low streaming whereas a region scoring high would have less differentiated cancer cells and more streams of cells. This scoring system was applied to an evenly balanced selection of vascularised and non-vascularised fields of view from H&Es of 5 human OSCC specimens, in a blinded analysis performed independently by two researchers **(Figure 7A, B)**. A higher score was obtained in vascularised areas compared to non-vascularised areas suggesting that, in a vascularised environment, OSCC have a less differentiated presentation and invade as thinner structures resembling streams of cells. To further investigate this association, we analysed an independent collection of H&Es from 49 human OSCC tumours. These H&Es were scored in a blinded analysis performed independently by two pathologists. Tumours were scored on vascular density, invasive front (cohesive vs discohesive) and invasive pattern (broad front vs multiple strands). Low vascular density was associated with cohesive invasion as broad sheets, whereas higher vascular density was associated with discohesive invasion as multiple strands **(Figure 7C-E)**. This further supports the conclusion that the streaming OSCC invasion observed in vascularised microfluidic chips is also associated with vascularised regions *in vivo*, in human OSCC specimens.

**Figure 7.**
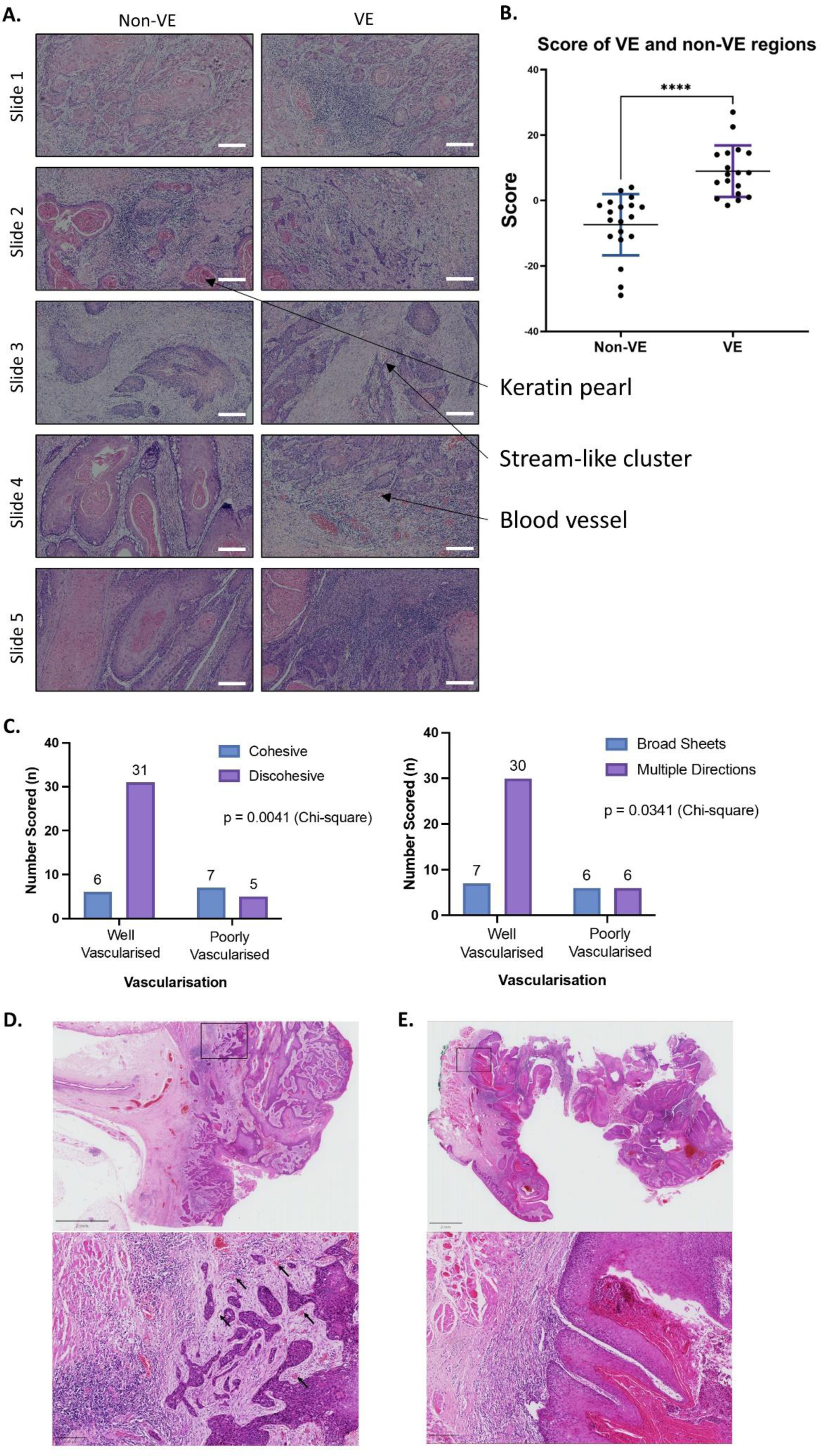
Association of streaming invasion with vascularised regions in human OSCC specimens. **A.** Fields of view of non-vascularised (non-VE) regions and vascularised (VE) regions from 5 human OSCC H&E stained tumour specimens. Examples of features included in the scoring system are indicated. Scale bars: 200 μm. **B.** Score of non-VE and VE regions in the H&E stained specimens. Mann-Whitney test. Data from 5 independent H&E specimens, with each dot representing a field of view within an H&E specimen. **C.** In feature scoring of H&E samples from 49 human OSCC tumours, association of vascularisation score with scoring for features of the invasive front is shown. In this analysis, each specimen receives a single binary score for each feature. **D.** An example tumour with a discohesive invasive front, multiple direction pattern of invasion and high vascular density. **E.** An example tumour with a cohesive invasive front, broad sheet pattern of invasion and low vascular density. For **(D)** and **(E)**, Top: Low magnification image, scale bar: 2 mm. Bottom: High magnification image (from the area indicated by the black box in the top image), with black arrows indicating examples of vasculature. Scale bar: 200 μm.

## Discussion

In this study, we have utilised a vascularised microfluidic chip to characterise the effect of local developing microvasculature on the invasive behaviour of OSCC. We observed that OSCC invasion is significantly reduced in the presence of a microvasculature. As the endothelial cells grow into the matrix, they remodel and compress the fibrin gel. This environment appears to be refractory to OSCC cell invasion; broad invading sheets of cells with single cell invasion at the leading edge, as seen in the control samples, is absent in this setting. Instead, the cancer cells utilise tracks within the matrix, made by single endothelial cells, to migrate into this environment as collective streams of cells **(Figure S5)**. The cells forming these collective OSCC streams express specific marker patterns consistent with a hybrid CSC phenotype that maintains cell-cell junctions (e-cadherin), downregulates epithelial markers (pan-keratin), expresses plasticity markers (CD24 and PDPN) and does not undergo full EMT (lacks Vimentin). The tip cells of these streams express high levels of CD44.

CD44 is a transmembrane glycoprotein that is a CSC marker in OSCC (Prince et al., 2007). Other studies have also demonstrated the presence of CD44 +ve cells at the edge of collective invasive masses (Gao et al., 2022, Wolf et al., 2020, Yang et al., 2019). For instance, Yang *et al*. (2019) showed that the invasive front of breast tumours exhibits a hybrid EMT phenotype, with high expression of CD44. Hence, CD44+ cells appear to play a significant role in leading collective invasion, and CD44 expression is associated with increased motility and matrix-adhesive characteristics.

Previous studies have shown that pre-existing tracks in the matrix can enable cancer cells to undergo stream-like invasion (Gaggioli et al., 2007, Kim et al., 2019, Vilchez Mercedes et al., 2021). The formation of these tracks has been associated with macrophages and fibroblasts. However, to our knowledge, invasive streaming behaviour has not previously been associated with tracks made by the developing vasculature.

Remodelling of matrix structure was only recorded proximally to the endothelial cells. When devices with a greater distance between the endothelial cells and OSCC cells were used, thicker fibrin fibres were only detected near to the endothelial cells and the stream-like phenotype of OSCC cells was absent. This is consistent with a mechanism whereby direct interaction between the endothelial cells and surrounding ECM is responsible for the observed remodelling. Endothelial cells have the ability to interact strongly with a fibrin ECM, notably with the expression of integrin αvβ3, recognised to interact directly with fibrin proteins (Feng et al., 2013). Once endothelial cells are activated in angiogenesis, tip cells grow through the fibrin and remodel it using proteases, including MT1-MMP (Lafleur et al., 2002). Through this process and interactions with the fibrin, sprouts and new vessels are formed. Previous studies have demonstrated that, during angiogenesis, sprouting endothelial cells remodel the surrounding matrix and increase its stiffness. This has been shown to occur only within 50 µm of sprouting endothelial cells (Juliar et al., 2018), a finding that is in agreement with our own data. This effect is driven by cell contractility driving matrix contraction, and is required for vascular tube formation (Rüdiger et al., 2020). Thus, the developing vasculature is guided into collective migration along defined channels. This is dependent on the ECM; in matrices which they cannot contract, such as collagen, HUVECs migrate as single cells and do not form a vasculature (Feng et al., 2013). We have now shown that this same guided approach is co-opted by OSCC CSCs for tissue invasion within a vascularised ECM.

The expression of CSC plasticity markers CD24 and PDPN suggests that the streaming OSCC cells may be capable of altering their phenotype during invasion, in response to environmental challenges. During collective invasion, the ability to loosen cell-cell junctions can allow cells to adapt to the extracellular space (Ilina et al., 2020). Thus, CSC plasticity may enable invasion plasticity, where cells adapt to the extracellular space to drive invasion through different spatial constrictions. This plasticity of invasive phenotype may be essential to enable streaming invasion under vascularised conditions. Notably, the non-transformed NTERT cell line was able to invade as a broad pushing front under control conditions, but lacked the ability to invade in the vascularised chips. Unlike the OSCC lines, they lacked invasion plasticity and were unable to utilise the tracks to undergo streaming invasion.

One surprising observation was the loss of Vimentin+ single cell invasion in the vascularised chips. Full EMT at the invasive front was instead replaced with collective invasion of a hybrid CSC. Whether the remodelled ECM is refractory to single cell invasion, or whether the endothelial cells inhibit full EMT in the OSCC cells through a different mechanism, remains to be explored. FACS of cells from 2D culture using hybrid CSC markers showed enrichment for streaming invasion, demonstrating that the hybrid CSCs responsible for streaming invasion are a pre-existing population and not a *de novo* adaptation to a vascularised environment. The ability to transition between different invasive phenotypes (broad front, single cell, streaming) through invasion plasticity, and thus adapt to overcome TME constraints, emerges as an important feature of CSC-driven tumour invasion.

The interaction of a tumour with its TME is an increasingly important avenue for therapeutic development, whereby the TME may be therapeutically targeted to modulate tumour behaviour. However, to achieve this we need a detailed understanding of tumour-TME interactions at a single cell level. Microfluidic metastasis-on-a-chip models are well placed to fulfil this aim, with their utility for precise imaging and real-time studies at single cell resolution within a complex cellular environment. In so doing, they also provide a step towards replacement of mouse models for TME studies. In combination with human tissue studies, they can deepen our understanding of tumour-TME interactions. Here, we have used such an approach to unravel the dynamic relationship between CSCs and the vasculature during OSCC invasion.

## Supporting information

Supplementary figures for Scemama et al

## Acknowledgements

We thank Liisa Blowes at the Blizard Institute CREATE lab and Luke Gammon at the Blizard Institute Phenotypic Screening Facility for technical assistance. Alice Scemama was supported by the National Centre for the 3Rs (NC3Rs) (NC/S001573/1). Sophia Lunetto was supported by the BBSRC LIDo DTP. Artysha Tailor was supported by The Medical College of Saint Bartholomew’s Hospital Trust. Leah Ambler was supported by Oracle Cancer Trust. Abigail Coetzee and Adrian Biddle were supported by the UK Medical Research Council (MRC) (MR/V009494/1). Adrian Biddle is a member of the Barts Centre for Squamous Cancer, funded by Barts Charity (G-002030).

## Methods

### Cell culture

The CA1 and LUC4 cell lines were both previously derived in our laboratory, from biopsies of OSCC of the floor of the mouth. The CA1 cell line was also previously retrovirally transduced with GFP to produce a GFP+ derivative of this cell line (Scemama et al., 2023). The NTERT cell line is a hTERT-immortalised epidermal keratinocyte cell line (Dickson et al., 2000). Cell culture was performed as previously described (Biddle et al., 2011). Detachment of OSCC cells and NTERTs from adherent surface was performed using 0.05% Trypsin-EDTA 1X (Sigma). Cells were cultured at 37°C with 5% CO2.

HUVECs (Lonza) and RFP-HUVECs (Angioproteomie) were cultured in EGM-2 medium (Promocell) until passage 6 and 8 respectively. Cells were detached using 0.05% Trypsin-EDTA 1X (Sigma). Medium was replaced every 48 hours. Cells were cultured at 37°C with 5% CO2.

Detailed protocols for cell culture are available in our recent method article (Scemama et al., 2023).

### Microfluidic device fabrication

The microfluidic devices were fabricated using photolithography, with the photomasks designed on AutoCAD®. A silicon wafer was spin-coated with SU8-2050 (Kayaku Advanced Materials), followed by a soft bake step. The photomask was placed on the wafer and illuminated with UV (45 mW/cm^2^) for 1 second. A post-exposure bake step followed, before immersing the wafer in PGMEA for the developing step and a final hard bake step was performed. PDMS was poured onto the wafer bearing the chip pattern and cross-linked by exposing it to 60-80°C for 1.5 hours. Devices were cleaned and plasma bonded to glass cover slips to seal the microchannels. Autoclaving of the microfluidic chips was performed before being used for cell culture experiments.

Detailed protocols for microfluidic device fabrication are available in our recent method article (Scemama et al., 2023).

### Cell culture in the microfluidic chips

A fibrin gel was prepared by combining equal parts fibrinogen from bovine plasma (10–15 mg/ml in PBS, Sigma) and thrombin from bovine plasma (5 U/ml in PBS, Sigma), and added in the middle channel of the microfluidic chip. For fluorescent particle experiments, fluorescent particles (Bangslabs UMGB003) were sterilised in 70% ethanol, washed twice with sterile PBS and then resuspended in 500 μl of sterile PBS. This suspension was used as the base for the thrombin solution for these experiments.

HUVECs were seeded the following day by adding 8 µl of a 5-7.5 million cells/ml cell suspension in one of the inlets. The devices were flipped at 90° for 30 minutes to allow the cells to adhere to the gel interface. EGM-2 medium supplemented with VEGF (50 ng/ ml, Recombinant Human VEGF, Peprotech 100-20A) was then added in all the inlets and an interstitial flow was created by having different volumes of medium in the inlets (90ul in the inlets of the HUVEC channel and 110ul in the inlets of the opposite channel). Medium was replaced every 24 hours. 2-3 days later, 8 µl of a 5 million cells/ml OSCC/NTERTs cell suspension was added in one of the inlets of the opposite side channel and allowed to adhere to the gel interface by flipping the device at 90° for 30 minutes. Fresh medium was added in the same way, and replaced every 24 hours. Chips were fixed at timepoints for imaging, using a 4% paraformaldehyde solution (Thermofisher) for 15 minutes, washed in PBS and then transferred to the imaging protocol.

Detailed protocols for cell culture in the microfluidic chip are available in our recent method article (Scemama et al., 2023).

### FACS sorting

Cells were detached using 0.05% Trypsin-EDTA 1X (Sigma) and stained with the following conjugated antibodies (1/100 dilution in PBS) for 15 minutes at 4°C: CD44 (PE, BD Biosciences 555479), EpCAM (FITC, Miltenyi Biotec 130-110-998) and PDPN (APC, Biolegend 337022). To exclude dead cells from the analysis, cells were stained with DAPI (1/2000 in sterile PBS from 1 mg/ml stock, Roche). Sorted cells were re-suspended in medium to reach a 2.5 million cells/ml concentration. 8 µl of this cell suspension was added in one of the inlets of the microfluidic chip, which was then flipped at 90° to allow the cells to adhere to the gel interface.

### Immunofluorescent staining and imaging

Chips were permeabilised with 0.1% Triton X-100 (Sigma-Aldrich) for 10 minutes. Samples to be stained with conjugated antibodies were first blocked with a 3% BSA solution in PBS (blocking buffer) for 3 hours at room temp. Conjugated antibodies, diluted in blocking buffer, were then added in all the inlets and left overnight at 4°C. Conjugated antibodies used for this work were as follow: Vimentin (Alexa Fluor 488, 1/500 dilution, Abcam ab195877), Cytokeratin (Alexa Fluor 647, 1/100 dilution, Biolegend 628604), CD44 (APC, 1/500 dilution, Miltenyi 130-113-331), EpCAM (Alexa Fluor 488, 1/100 dilution, Abcam ab237395), PDPN (APC, 1/500 dilution, Biolegend 337022), CD24 (APC, 1/200 dilution, Biolegend 311117) and E-cadherin (Alexa Fluor 647, 1/500 dilution, BD Pharmingen 563571). A non-conjugated Ki67 antibody (1/500 dilution, Abcam ab833) was used with an Alexa Fluor 555 goat anti-rabbit secondary antibody (1/1000 dilution, Invitrogen A21429). Chips stained with CellMask (1/50,000 dilution in PBS, Thermo Fisher) were permeabilised with 0.1% Triton X-100 (Sigma-Aldrich) for 10 minutes but not blocked. The CellMask stain was left in the device for 2 hours. If stained with phalloidin (1/500, Scientific Laboratory Supplies P1951) and DAPI (1/1000 in PBS from 1 mg/ml stocks, Roche 10236276001), or just DAPI, these stains were mixed with the blocking buffer directly and added in each microfluidic chip inlet for one hour, at room temp, after antibody staining. Chips were then washed twice with PBS and left in PBS at 4°C until imaged. Imaging was performed using the IN Cell Analyzer 6000 (GE Healthcare). Five to six fields of view, using the 20× magnification, were taken to cover the entire middle channel for each chip. Twenty stacks (4 μm thick each) were taken per field of view. Stacks were maximum intensity projected. FIJI, ICY and Developer Toolbox (GE Healthcare) were used for image analysis. Detailed protocols for image analysis are available in our recent method article (Scemama et al., 2023).

### Matrix structure imaging

The Zeiss 880 Laser Scanning Confocal Microscope with Fast Airyscan and Multiphoton microscope (inverted, Zeiss 880 Confocal microscope) was used to image the matrix structure via reflectance confocal imaging. A 4.5 μm layer was imaged. The parameters selected on the ZEISS ZEN software were as follows: interval: 0.5, slices: 10, optimal: 0.33 μm. The matrix was imaged using the 488 channel. Stacks were maximum intensity projected. Developer Toolbox (GE Healthcare) were used for image analysis.

### FITC-Dextran assay

Medium was aspirated from the inlets of the microfluidic chip. FITC-Dextran 70 (FD70S, Sigma-Aldrich) was mixed with EGM-2 medium to reach a final concentration of 50 μg/ml. 60 μl of this mixture was added in the HUVEC side channel of the chip and max intensity projections were taken every 5 minutes for 30 minutes using the IN Cell Analyzer 6000 (GE Healthcare) microscope.

### Analysis of human OSCC H&E specimens

Human OSCC archival FFPE specimens were selected from the diagnostic archive at Barts Health NHS Trust by a consultant pathologist, with full ethical approval (REC reference: 18/WM/0326). H&E slides were prepared by the Blizard Institute pathology core. Scoring was performed blinded, by two researchers for the initial analysis of 5 specimens and by two pathologists for the subsequent analysis of 49 specimens. Scoring was performed independently by each individual, and scores were then averaged. For the analysis of 49 tumours, an agreement score of 85% between the two scorers was recorded. The tumours included in the analysis were balanced on T and N stage (TNM 7^th^ edition). For the initial analysis of 5 specimens, 3-4 each of vascularised and non-vascularised fields of view were selected from each tumour and scored separately. For the subsequent analysis of 49 specimens, the entire tumour slide was assessed and a single binary score for each feature was assigned per specimen.

### Statistical analysis

All data are based on at least three independent experimental repeats and with each datapoint representing a single microfluidic chip, unless specified in the legend. Statistical analysis was performed using the GraphPad Prism software, with Welch’s t-test used for statistical comparison unless specified in the legend. Statistical significance was considered achieved for p < 0.05, with * referring to p < 0.05, ** to p < 0.01, *** to p < 0.001 and **** to p < 0.0001. Bar graphs are presented as mean with standard deviation.

## References

Anderson NM, Simon MC. The tumor microenvironment. Curr Biol 2020;30(16):R921–r5.

Biddle A, Gammon L, Liang X, Costea DE, Mackenzie IC. Phenotypic Plasticity Determines Cancer Stem Cell Therapeutic Resistance in Oral Squamous Cell Carcinoma. EBioMedicine 2016;4:138–45.

Biddle A, Liang X, Gammon L, Fazil B, Harper LJ, Emich H, et al. Cancer stem cells in squamous cell carcinoma switch between two distinct phenotypes that are preferentially migratory or proliferative. Cancer Res 2011;71(15):5317–26.

Chaffer CL, San Juan BP, Lim E, Weinberg RA. EMT, cell plasticity and metastasis. Cancer and Metastasis Reviews 2016;35(4):645–54.

Chen MB, Whisler JA, Jeon JS, Kamm RD. Mechanisms of tumor cell extravasation in an in vitro microvascular network platform. Integr Biol (Camb) 2013;5(10):1262–71.

Clarke MF, Dick JE, Dirks PB, Eaves CJ, Jamieson CH, Jones DL, et al. Cancer stem cells--perspectives on current status and future directions: AACR Workshop on cancer stem cells. Cancer Res 2006;66(19):9339–44.

Dickson MA, Hahn WC, Ino Y, Ronfard V, Wu JY, Weinberg RA, et al. Human keratinocytes that express hTERT and also bypass a p16(INK4a)-enforced mechanism that limits life span become immortal yet retain normal growth and differentiation characteristics. Mol Cell Biol 2000;20(4):1436–47.

Driessens G, Beck B, Caauwe A, Simons BD, Blanpain C. Defining the mode of tumour growth by clonal analysis. Nature 2012;488(7412):527-30.

Ehrbar M, Sala A, Lienemann P, Ranga A, Mosiewicz K, Bittermann A, et al. Elucidating the role of matrix stiffness in 3D cell migration and remodeling. Biophys J 2011;100(2):284–93.

Feng X, Tonnesen MG, Mousa SA, Clark RA. Fibrin and collagen differentially but synergistically regulate sprout angiogenesis of human dermal microvascular endothelial cells in 3-dimensional matrix. Int J Cell Biol 2013;2013:231279.

Ferguson LP, Diaz E, Reya T. The Role of the Microenvironment and Immune System in Regulating Stem Cell Fate in Cancer. Trends Cancer 2021;7(7):624–34.

Friedl P, Alexander S. Cancer invasion and the microenvironment: plasticity and reciprocity. Cell 2011;147(5):992–1009.

Friedl P, Locker J, Sahai E, Segall JE. Classifying collective cancer cell invasion. Nature Cell Biology 2012;14(8):777–83.

Gaggioli C, Hooper S, Hidalgo-Carcedo C, Grosse R, Marshall JF, Harrington K, et al. Fibroblast-led collective invasion of carcinoma cells with differing roles for RhoGTPases in leading and following cells. Nat Cell Biol 2007;9(12):1392–400.

Gao F, Zhang G, Liu Y, He Y, Sheng Y, Sun X, et al. Activation of CD44 signaling in leader cells induced by tumor-associated macrophages drives collective detachment in luminal breast carcinomas. Cell Death Dis 2022;13(6):540.

Ilina O, Gritsenko PG, Syga S, Lippoldt J, La Porta CAM, Chepizhko O, et al. Cell-cell adhesion and 3D matrix confinement determine jamming transitions in breast cancer invasion. Nat Cell Biol 2020;22(9):1103–15.

Jensen DH, Dabelsteen E, Specht L, Fiehn AM, Therkildsen MH, Jønson L, et al. Molecular profiling of tumour budding implicates TGFβ-mediated epithelial-mesenchymal transition as a therapeutic target in oral squamous cell carcinoma. J Pathol 2015;236(4):505–16.

Juliar BA, Keating MT, Kong YP, Botvinick EL, Putnam AJ. Sprouting angiogenesis induces significant mechanical heterogeneities and ECM stiffening across length scales in fibrin hydrogels. Biomaterials 2018;162:99–108.

Kim H, Chung H, Kim J, Choi DH, Shin Y, Kang YG, et al. Macrophages-Triggered Sequential Remodeling of Endothelium-Interstitial Matrix to Form Pre-Metastatic Niche in Microfluidic Tumor Microenvironment. Adv Sci (Weinh) 2019;6(11):1900195.

Kim S, Lee H, Chung M, Jeon NL. Engineering of functional, perfusable 3D microvascular networks on a chip. Lab Chip 2013;13(8):1489–500.

Kniazeva E, Weidling JW, Singh R, Botvinick EL, Digman MA, Gratton E, et al. Quantification of local matrix deformations and mechanical properties during capillary morphogenesis in 3D. Integr Biol (Camb) 2012;4(4):431–9.

Kreso A, O’Brien CA, van Galen P, Gan OI, Notta F, Brown AM, et al. Variable clonal repopulation dynamics influence chemotherapy response in colorectal cancer. Science 2013;339(6119):543–8.

Krishnamurthy S, Dong Z, Vodopyanov D, Imai A, Helman JI, Prince ME, et al. Endothelial cell-initiated signaling promotes the survival and self-renewal of cancer stem cells. Cancer Res 2010;70(23):9969–78.

Lafleur MA, Handsley MM, Knäuper V, Murphy G, Edwards DR. Endothelial tubulogenesis within fibrin gels specifically requires the activity of membrane-type-matrix metalloproteinases (MT-MMPs). Journal of cell science 2002;115(Pt 17):3427–38.

Lawson DA, Bhakta NR, Kessenbrock K, Prummel KD, Yu Y, Takai K, et al. Single-cell analysis reveals a stem-cell program in human metastatic breast cancer cells. Nature 2015;526(7571):131–5.

Lee H, Park W, Ryu H, Jeon NL. A microfluidic platform for quantitative analysis of cancer angiogenesis and intravasation. Biomicrofluidics 2014;8(5):054102.

Lee JH, Kim SK, Khawar IA, Jeong SY, Chung S, Kuh HJ. Microfluidic co-culture of pancreatic tumor spheroids with stellate cells as a novel 3D model for investigation of stroma-mediated cell motility and drug resistance. J Exp Clin Cancer Res 2018;37(1):4.

Mehlen P, Puisieux A. Metastasis: a question of life or death. Nat Rev Cancer 2006;6(6):449–58.

Oshimori N, Oristian D, Fuchs E. TGF-β promotes heterogeneity and drug resistance in squamous cell carcinoma. Cell 2015;160(5):963–76.

Pandya P, Orgaz JL, Sanz-Moreno V. Modes of invasion during tumour dissemination. Mol Oncol 2017;11(1):5–27.

Pastushenko I, Mauri F, Song Y, de Cock F, Meeusen B, Swedlund B, et al. Fat1 deletion promotes hybrid EMT state, tumour stemness and metastasis. Nature 2021;589(7842):448–55.

Prince ME, Sivanandan R, Kaczorowski A, Wolf GT, Kaplan MJ, Dalerba P, et al. Identification of a subpopulation of cells with cancer stem cell properties in head and neck squamous cell carcinoma. Proc Natl Acad Sci U S A 2007;104(3):973–8.

Puram SV, Tirosh I, Parikh AS, Patel AP, Yizhak K, Gillespie S, et al. Single-Cell Transcriptomic Analysis of Primary and Metastatic Tumor Ecosystems in Head and Neck Cancer. Cell 2017;171(7):1611–24 e24.

Rizvi I, Gurkan UA, Tasoglu S, Alagic N, Celli JP, Mensah LB, et al. Flow induces epithelial-mesenchymal transition, cellular heterogeneity and biomarker modulation in 3D ovarian cancer nodules. Proc Natl Acad Sci U S A 2013;110(22):E1974–83.

Rüdiger D, Kick K, Goychuk A, Vollmar AM, Frey E, Zahler S. Cell-Based Strain Remodeling of a Nonfibrous Matrix as an Organizing Principle for Vasculogenesis. Cell Rep 2020;32(6):108015.

Sano D, Myers JN. Metastasis of squamous cell carcinoma of the oral tongue. Cancer metastasis reviews 2007;26(3-4):645–62.

Scemama A, Lunetto S, Biddle A. Highlight: microfluidic devices for cancer metastasis studies. In vitro models 2022;1(6):399–403.

Scemama A, Tailor A, Di Cio S, Dibble M, Gautrot J, Biddle A. Development of an in vitro microfluidic model to study the role of microenvironmental cells in oral cancer metastasis [version 1; peer review: awaiting peer review]. F1000Research 2023;12(439).

Taniguchi S, Elhance A, Van Duzer A, Kumar S, Leitenberger JJ, Oshimori N. Tumor-initiating cells establish an IL-33-TGF-beta niche signaling loop to promote cancer progression. Science 2020;369(6501).

Thiery JP. Epithelial-mesenchymal transitions in tumour progression. Nat Rev Cancer 2002;2(6):442–54.

Tsai JH, Donaher JL, Murphy DA, Chau S, Yang J. Spatiotemporal regulation of epithelial-mesenchymal transition is essential for squamous cell carcinoma metastasis. Cancer cell 2012;22(6):725–36.

van Roy F, Berx G. The cell-cell adhesion molecule E-cadherin. Cell Mol Life Sci 2008;65(23):3756–88.

Vilchez Mercedes SA, Bocci F, Levine H, Onuchic JN, Jolly MK, Wong PK. Decoding leader cells in collective cancer invasion. Nature Reviews Cancer 2021;21(9):592–604.

Wolf KJ, Shukla P, Springer K, Lee S, Coombes JD, Choy CJ, et al. A mode of cell adhesion and migration facilitated by CD44-dependent microtentacles. Proc Natl Acad Sci U S A 2020;117(21):11432–43.

Wu K, Wei J, Liu Z, Yu B, Yang X, Zhang C, et al. Can pattern and depth of invasion predict lymph node relapse and prognosis in tongue squamous cell carcinoma. BMC Cancer 2019;19(1):714.

Yang C, Cao M, Liu Y, He Y, Du Y, Zhang G, et al. Inducible formation of leader cells driven by CD44 switching gives rise to collective invasion and metastases in luminal breast carcinomas. Oncogene 2019;38(46):7113–32.

Yang MH, Wu MZ, Chiou SH, Chen PM, Chang SY, Liu CJ, et al. Direct regulation of TWIST by HIF-1alpha promotes metastasis. Nat Cell Biol 2008;10(3):295–305.

Youssef G, Gammon L, Ambler L, Lunetto S, Scemama A, Cottom H, et al. Disseminating cells in human oral tumours possess an EMT cancer stem cell marker profile that is predictive of metastasis in image-based machine learning. eLife 2023;12:e90298.

Zaman MH, Trapani LM, Sieminski AL, Mackellar D, Gong H, Kamm RD, et al. Migration of tumor cells in 3D matrices is governed by matrix stiffness along with cell-matrix adhesion and proteolysis. Proc Natl Acad Sci U S A 2006;103(29):10889–94.

Zervantonakis IK, Hughes-Alford SK, Charest JL, Condeelis JS, Gertler FB, Kamm RD. Three-dimensional microfluidic model for tumor cell intravasation and endothelial barrier function. Proc Natl Acad Sci U S A 2012;109(34):13515–20.

