## Supplementary figures for Scemama et al for "Hybrid cancer stem cells utilise vascular tracks for collective streaming invasion in a metastasis-on-a-chip device"

**Figure S1**

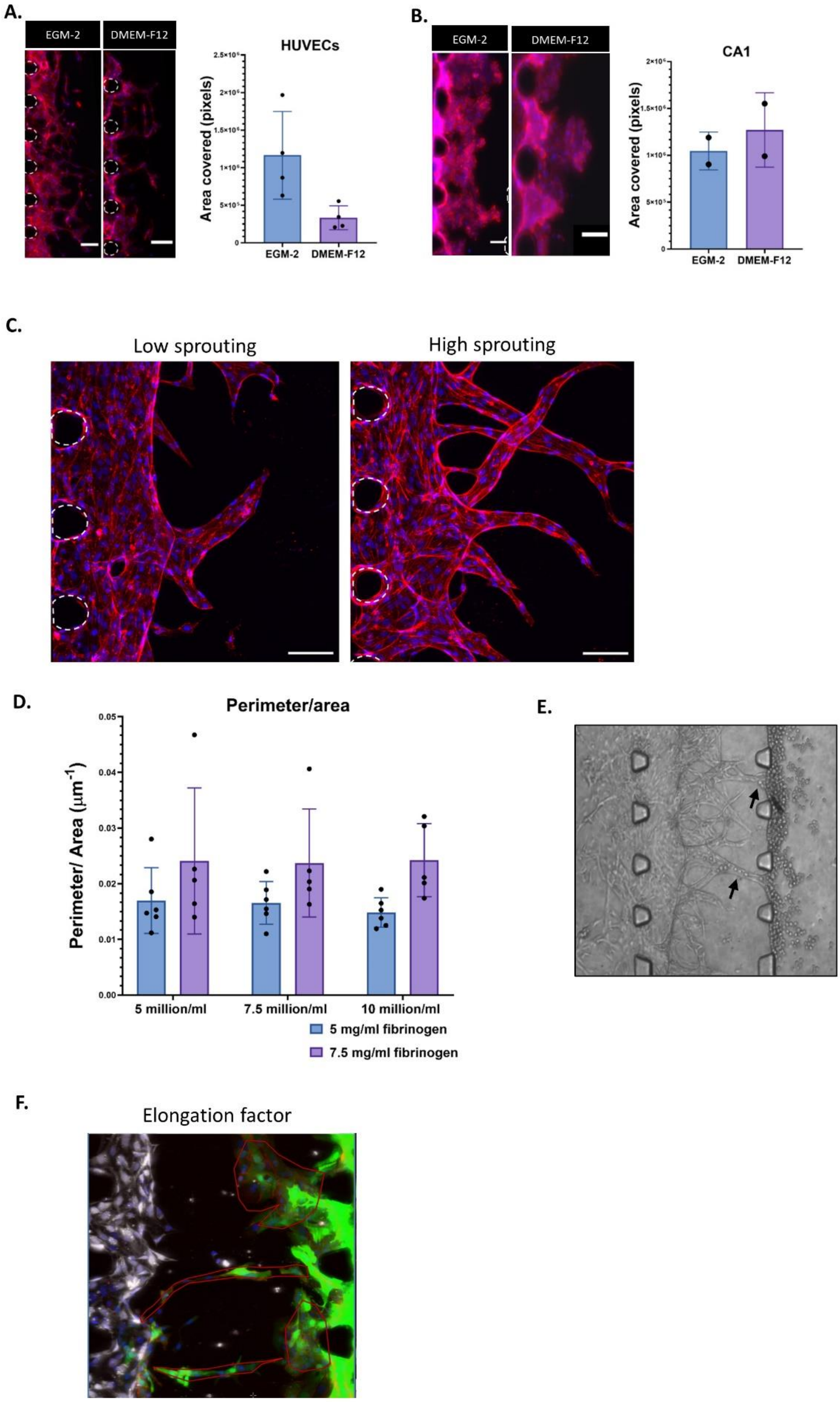

A.
EGM-2
DMEM-F12
HUVECs
Area covered (pixels)
EGM-2
DMEM-F12
B.
EGM-2
DMEM-F12
CA1
Area covered (pixels)
EGM-2
DMEM-F12
C.
Low sprouting
High sprouting
D.
Perimeter/area
Perimeter/ Area (μm$^{-1}$)
0.05
0.04
0.03
0.02
0.01
0.00
5 million/ml
7.5 million/ml
10 million/ml
5 mg/ml fibrinogen
7.5 mg/ml fibrinogen
E.
F.
Elongation factor

**Figure S1 - Optimisation of co-culture in the microfluidic chip**

**A, B: The choice of medium for co-culture.** OSCC cells and HUVECs normally grow in different growth medium, with DMEM-F12 and EGM-2 medium used respectively. We therefore tested which growth medium combination would enable culture of these two cell types in the microfluidic chip simultaneously. HUVEC growth into the central channel was retarded by the presence of DMEM-F12 in the opposite side channel **(A)**. Red, Phalloidin; Blue, DAPI. Posts are highlighted in white (dashed line). Scale bars: 100 µm. Data from 2 independent experiments, with each dot representing a single microfluidic chip. Conversely, growth of the CA1 OSCC cell line into the central channel was not significantly affected by the presence of EGM-2 medium in the opposite side channel **(B)**. Red, Phalloidin; Blue, DAPI. Posts are highlighted in white (dashed line). Scale bars: 100 µm. Data from 2 independent experiments, with each dot representing the average of 1-5 microfluidic chips in each experiment. We therefore elected to use EGM-2 medium for co-culture in the microfluidic chip, to ensure appropriate growth of the endothelial cells.

**C-E: Production of a lumenised vasculature.** Quantification of the level of vascular sprouting at day 11 after HUVEC addition **(C)** by dividing the vascular perimeter by area **(D)** showed that adding 5 – 7.5 million HUVEC per ml and using a final fibrinogen concentration of 5 – 7.5 mg/ml produced reliable vascular sprouting in the device. Red, Phalloidin; Blue, DAPI. Posts are highlighted in white (dashed line). Scale bars: 100 µm. Moreover, once the vasculature reaches all the way to the opposite side channel (day 11), cancer cells added at this point can flow into it (indicated by black arrows) **(E)**. This demonstrates that the vasculature is lumenised.

**F: Quantification of tumour cell invasion.** Calculation of area covered by the cells using Developer Toolbox, and calculation of the length of the cell front using FIJI, are covered in our accompanying methods paper (Scemama et al., 2023). Calculation of elongation factor of invasive masses using ICY was performed by drawing a region of interest around each invasive mass and then calculating the *elongation factor* value for each region **(F)**. The researcher was blinded when performing this analysis.

**Figure S2**

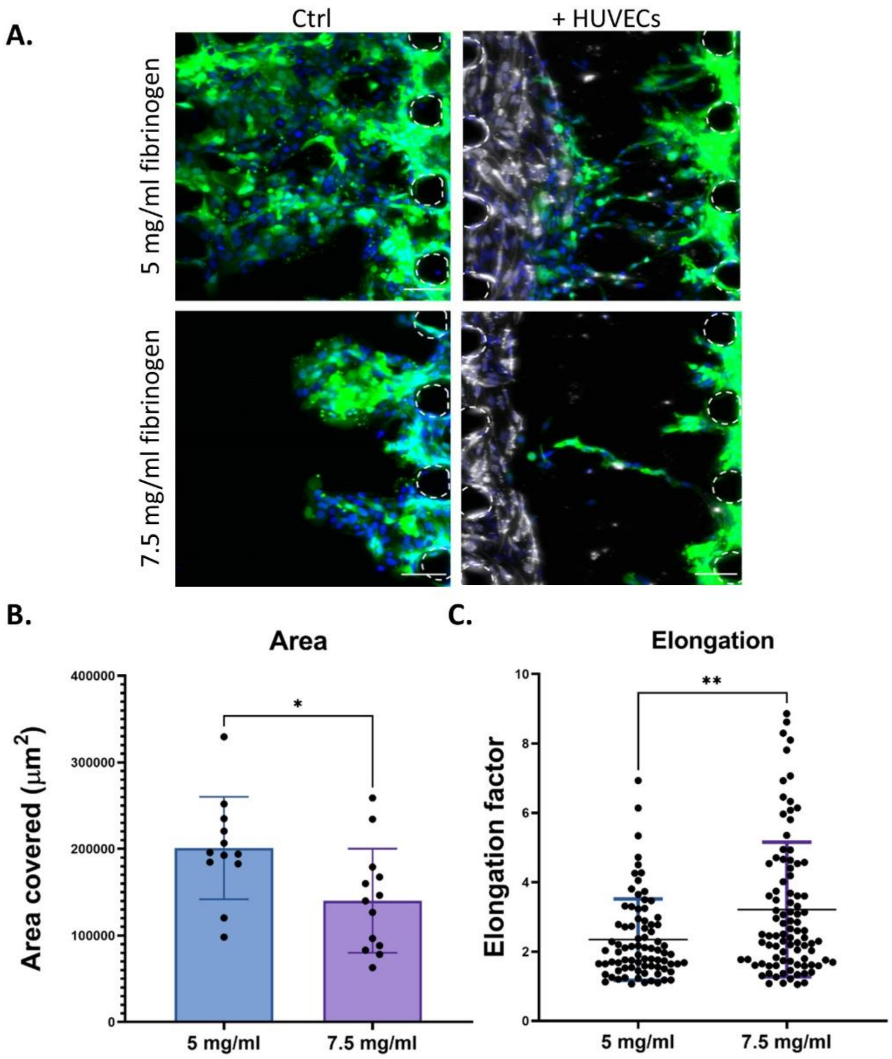


**Figure S2 – Increasing fibrinogen concentration results in reduced OSCC invasion and increased invasive front elongation**

**A:** CA1 cells only (Ctrl) and CA1 cells with HUVECs in a 5 mg/ml or 7.5 mg/ml fibrinogen gel at day 10-11. Green, GFP-CA1; Grey, RFP-HUVECs; Blue, DAPI. Posts are highlighted in white (dashed line). Scale bars: 100 μm.

**B:** Area covered by the CA1 cells in the 5 mg/ml and 7.5 mg/ml fibrinogen gels + HUVECs.

**C:** Elongation factor of the invasive streams of cells in the 5 mg/ml and 7.5 mg/ml fibrinogen gels + HUVECs. Data from 4 independent experiments, with each dot representing a single invasive mass within a microfluidic chip. Mann-Whitney test.

### Figure S3

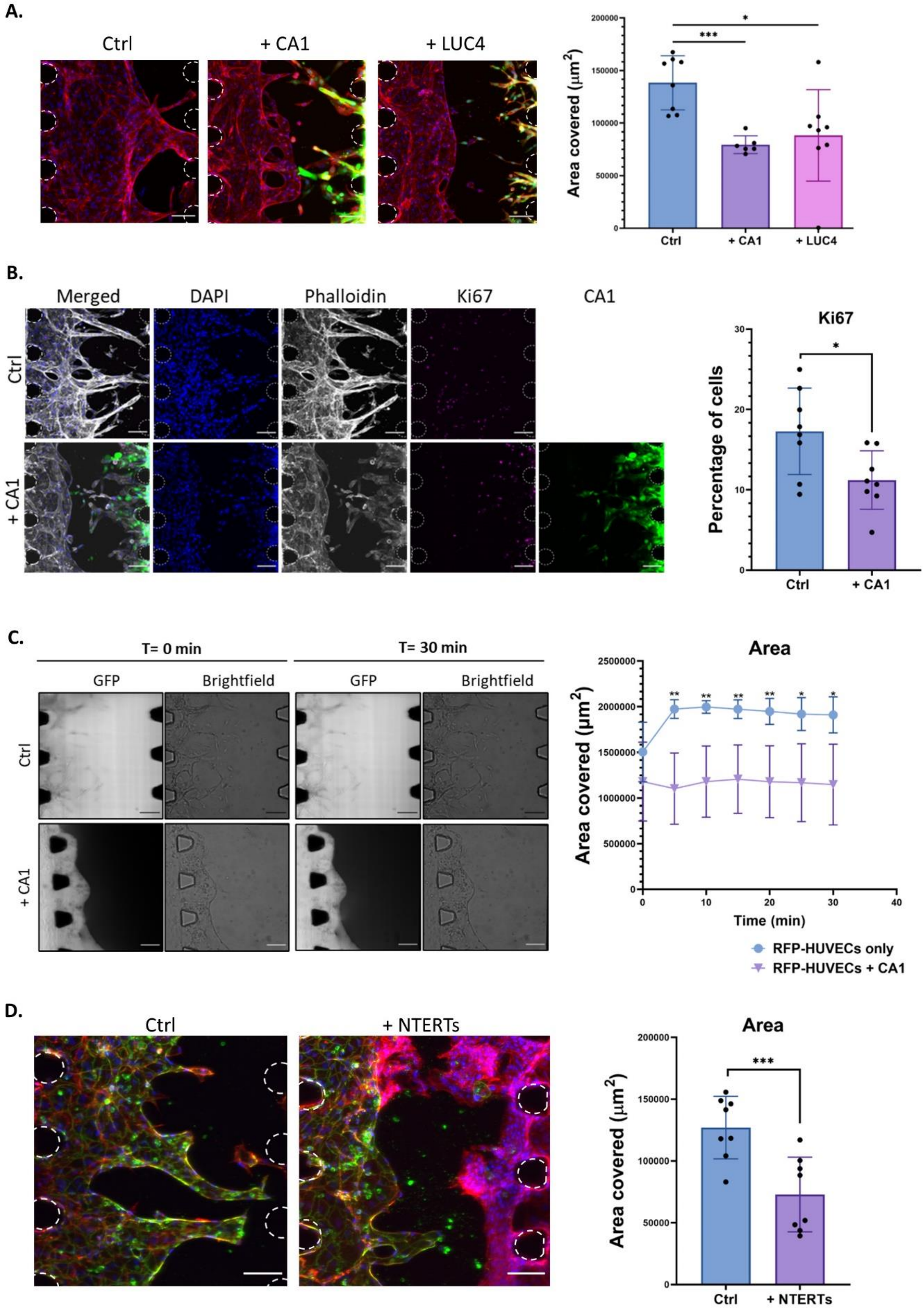
A.
Ctrl
+ CA1
+ LUC4
Area covered (μm²)
200000
150000
100000
50000
0
***
*
Ctrl
+ CA1
+ LUC4
B.
Merged
DAPI
Phalloidin
Ki67
CA1
Ctrl
+ CA1
Ki67
Percentage of cells
30
20
10
0
*
Ctrl
+ CA1
C.
T= 0 min
T= 30 min
GFP
Brightfield
GFP
Brightfield
Ctrl
+ CA1
Area
Area covered (μm²)
2500000
2000000
1500000
1000000
500000
0
**
**
**
**
*
*
0
10
20
30
40
Time (min)
RFP-HUVECs only
RFP-HUVECs + CA1
D.
Ctrl
+ NTERTs
Area
Area covered (μm²)
200000
150000
100000
50000
0
***
Ctrl
+ NTERTs

**Figure S3 – Behaviour of the microvasculature in response to OSCC cells**

**A: Vascular regression in the presence of OSCC cells.** Left: HUVECs only (Ctrl) and HUVECs with CA1 or LUC4 cells in the fibrin gel at day 11. Green, GFP-CA1/LUC4; Red, Phalloidin; Blue, DAPI. Posts are highlighted in white (dashed line). Scale bars: 100 μm. Right: Area covered by the HUVECs in the control (Ctrl) samples and samples with CA1 or LUC4 cells.

**B: Reduced Ki67 expression in HUVECs in the presence of CA1 cells.** Left: HUVECs only (Ctrl) and HUVECs with CA1 cells in the fibrin gel at day 11, stained with Ki67. Green, GFP-CA1; Grey, Phalloidin; Magenta, Ki67; Blue, DAPI. Posts are highlighted in white (dashed line). Scale bars: 100 μm. Right: Percentage of HUVECs expressing Ki67 in the control (Ctrl) samples and the samples with CA1 cells.

**C: Reduced FITC-Dextran permeability of HUVECs in the presence of CA1 cells.** Left: HUVECs only (Ctrl) and HUVECs with CA1 cells in the fibrin gel at day 11 with FITC-Dextran at T= 0 minutes and T= 30 minutes. Grey, FITC-Dextran. Scale bars: 100 μm. Right: Area covered by the FITC-Dextran over time in the control (Ctrl) samples and samples with CA1 cells. Multiple unpaired t-test.

**D: Vascular regression in the presence of NTERTs.** Left: HUVECs only (Ctrl) and HUVECs with NTERTs in the fibrin gel at day 11. Green, CD31; Red, Phalloidin; Blue, DAPI. Posts are highlighted in white (dashed line). Scale bars: 100 μm. Right: Area covered by the HUVECs in the control (Ctrl) samples and samples with NTERTs.

**Figure S4**

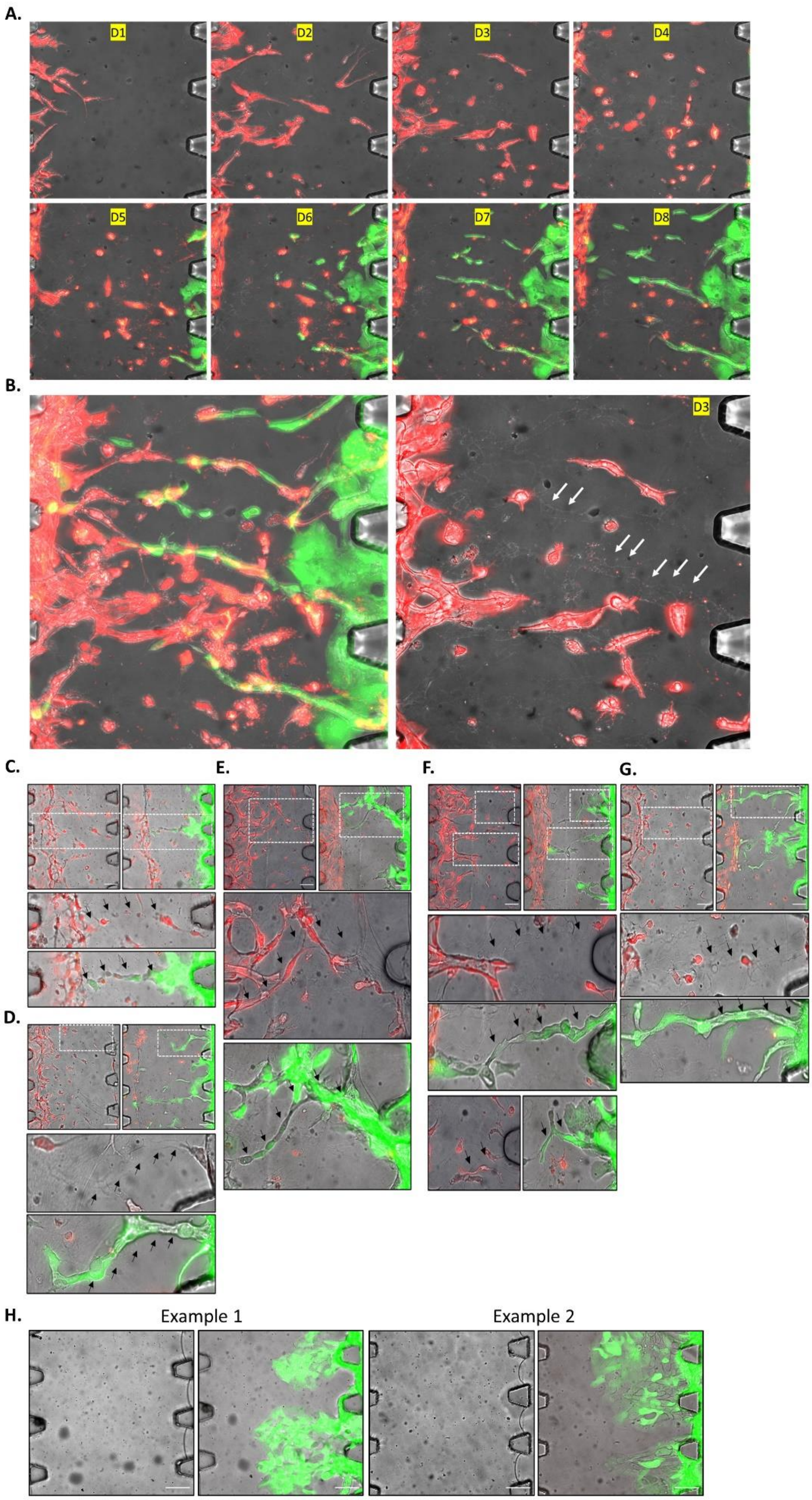

**Figure S4 – Further examples of the utilization of HUVEC-generated tracks for OSCC streaming invasion**

**A:** Daily timepoint analysis at days 1 to 8 in a vascularised chip, with HUVECs (Red; RFP) added at day 1 (D1) and CA1 OSCC cells (Green; GFP) added at day 4 (D4).

**B:** Left: Overlay projection of all timepoints from A, showing the CA1 OSCC cells (Green; GFP) taking the same route through the matrix as the HUVECs (Red; RFP). Right: The day 4 timepoint image with the HUVEC-produced tracks denoted with white arrows. A single RFP+ HUVEC can be seen at the head of a track, having progressed the whole way across the central channel.

**C-G:** Examples of the tracks generated by the HUVECs, showing the same field of view before addition of CA1 OSCC cells (left image, and top inset) and after CA1 addition (right image, and bottom inset). Black arrows denote the tracks produced by the HUVECs (Red; RFP), which are then utilised by the OSCC cells (Green; GFP) for invasion. Single RFP+ HUVECs can be seen at the head of each track before CA1 addition, having progressed the whole way across the central channel. Scale bars: 100 µm.

**H:** Control chips, without a vasculature, showing the same field of view before addition of CA1 OSCC cells (left image) and after CA1 addition (right image). Without a vasculature, no tracks are present and the CA1 OSCC cells (Green; GFP) invade across a broad front.

**Figure S5**

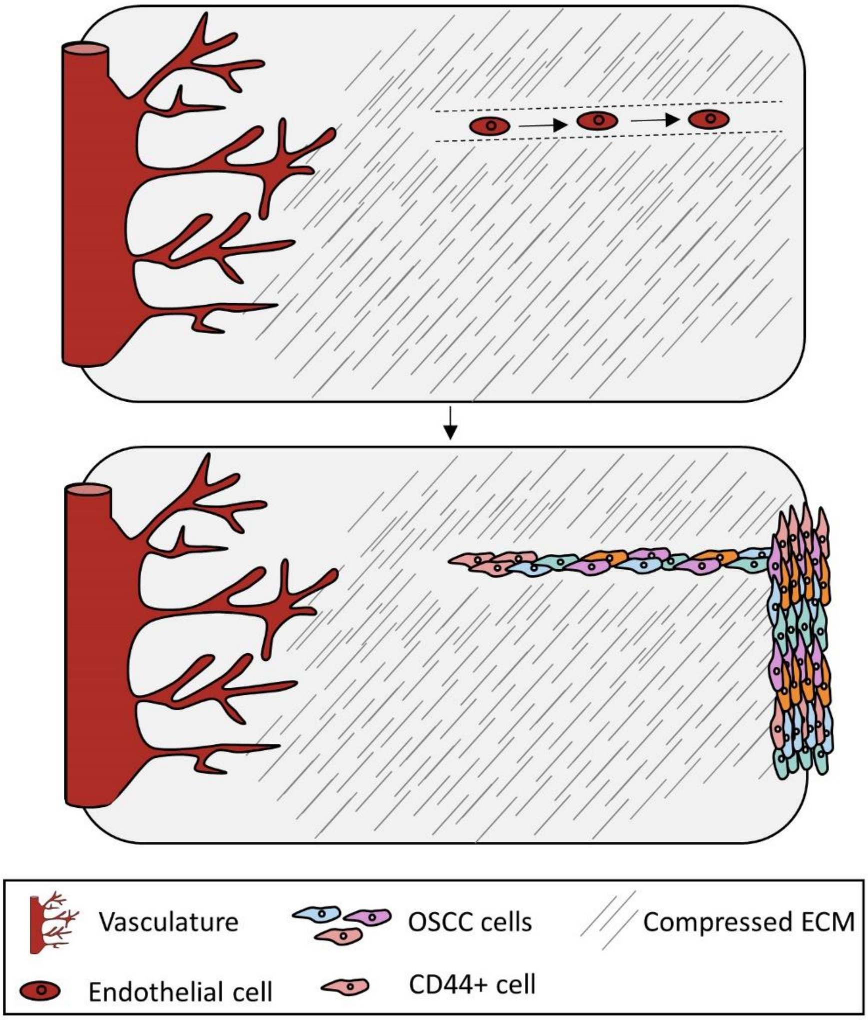


**Figure S5 – Summary of the effect of the developing microvasculature on tumour cell invasion.**

The developing microvasculature causes compression of the extracellular matrix in its vicinity, making it refractory to tumour cell invasion. Concurrently, individual vascular endothelial cells produce tracks in the matrix which guide vascular development. Upon addition of tumour cells, the vasculature regresses and leaves empty tracks within the compressed matrix. The tumour cells cannot invade into the compressed matrix, but instead are able to use the abandon vascular tracks to invade. The mode of tumour cell invasion is also altered, with the formation of collective invasive streams. These downregulate epithelial keratins and the EMT marker Vimentin. However, they upregulate CSC markers including CD44.
